# Structural origins of protein conformational entropy

**DOI:** 10.1101/2021.02.12.430981

**Authors:** José A. Caro, Kathleen G. Valentine, A. Joshua Wand

## Abstract

The thermodynamics of molecular recognition by proteins is a central determinant of complex biochemistry. For over a half-century detailed cryogenic structures have provided deep insight into the energetic contributions to ligand binding by proteins^1^. More recently, a dynamical proxy based on NMR-relaxation methods has revealed an unexpected richness in the contributions of conformational entropy to the thermodynamics of ligand binding^2,3,4,5^. There remains, however, a discomforting absence of an understanding of the structural origins of fast internal motion and the conformational entropy that this motion represents. Here we report the pressure-dependence of fast internal motion within the ribonuclease barnase and its complex with the protein barstar. Distinctive clustering of the pressure sensitivity correlates with the presence of small packing defects or voids surrounding affected side chains. Prompted by this observation, we performed an analysis of the voids surrounding over 2,500 methyl-bearing side chains having experimentally determined order parameters. We find that changes in unoccupied volume as small as a single water molecule surrounding buried side chains greatly affects motion on the subnanosecond timescale. The discovered relationship begins to permit construction of a united view of the relationship between changes in the internal energy, as exposed by detailed structural analysis, and the conformational entropy, as represented by fast internal motion, in the thermodynamics of protein function.

## Body

The change in the Gibbs free energy underlying molecular recognition and other complex protein functions such as allosteric regulation has, in principle, contributions from both entropy and enthalpy. The latter is comprised of the internal energy and a pressure-volume work term. Detailed analysis of static low-temperature structural models has historically provided great insight into the internal energy and has promoted significant advances in understanding protein functions such as ligand binding through simulation and theory^6^. Nevertheless, the origins of protein conformational entropy and its contribution to functions such as allostery remains less well defined^5^. Measurement of equilibrium fluctuations offers a powerful way to describe transitions between and occupancy of states that cannot be observed with classical methods of structural biology and NMR relaxation has proven particularly useful in this regard^4,7,8^. Over the past two decades, numerous studies of fast internal side chain motion by NMR methods, particularly that of methyl-bearing amino acids, have revealed an unexpected complexity without distinguishing structural correlates^9^.

Here we take advantage of the fact that the Gibbs free energy change associated with a change in state contains a pressure-volume work term. Volume changes represent the natural variable and application of pressure can illuminate otherwise unobservable details of the thermodynamics of protein functions such as ligand binding and allostery. Protein molecules respond, both dynamically and structurally, to pressure in a complicated way. Pressure can compress proteins^10,11,12^, remodel active sites^12^ and facilitate excursions to higher lying^13,14^, locally unfolded^15,16,17,18^ or globally unfolded states^19,20^, thereby revealing various aspects of the ensemble nature of proteins. Here, we use high pressure NMR^21^ relaxation to probe fast internal motion of methyl-bearing side chains in the small enzyme barnase and use this motion as a proxy for conformational entropy (S_conf_).

Barnase-barstar is one of the strongest protein-protein interactions known in biology. Its fM affinity derives from large enthalpic contributions at the interface^22^. Individual entropic contributions sum to yield a negligible (∼zero) contribution to the binding free energy^22^. At ambient pressure, we find that complexation is accompanied by an overall rigidification of the methyl-bearing side chains of barnase (Fig. 1; Supplementary Table 1). We quantify the disorder of the methyl symmetry axis in terms of the Lipari-Szabo squared generalized order parameter (O^2^_axis_)^23^ obtained using deuterium NMR relaxation methods^24^. The O^2^_axis_ can range from a value of one, corresponding to complete rigidity within the molecular frame, to zero, which effectively corresponds to isotropic disorder. Importantly, only motion faster than the overall molecular reorientation of the protein contributes to O^2^axis. The dynamical proxy for conformational entropy^4^ indicates that the overall rigidification corresponds to an unfavorable contribution (ΔS_conf_) to the binding free energy amounting to +11.7 ± 1.2 kJ/mol at room temperature.

**Fig. 1.**
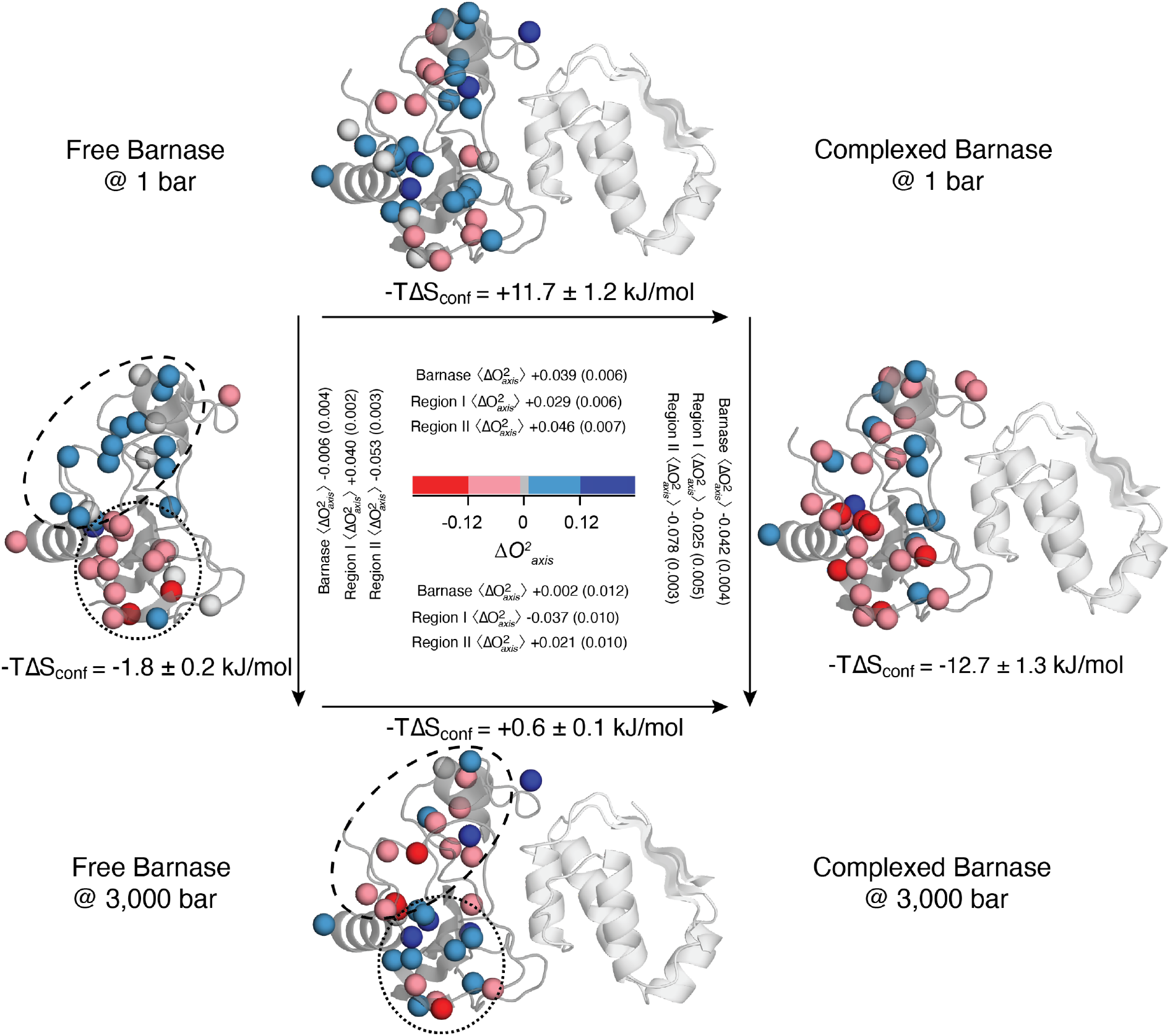
Dynamic response of barnase to high pressure and to binding barstar. The locations of reporter methyl groups are indicated by spheres on the ribbon-diagram of barnase (PDB 1B2X & 1B27) and color coded according to the 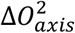 where red and blue correspond to decreases and increases in 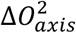 upon the indicated change of state, respectively. The inset summarizes the average change of 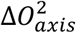 values in barnase as a whole and within the two regions indicated by the dashed lines, region I as dashed lines and region II as dotted lines (see text for details). Values in parentheses are the variance of changes within the indicated group and not error. Contributions of conformational entropy of the indicated state interconversion were obtained using the dynamical proxy^4^ and included all methyl probes. See Supplementary Table 1 for further details. [2 columns]

Application of high hydrostatic pressure on free barnase yields an unexpected clustering of changes in motion (ΔO^2^_axis_) into two spatial regions, one that rigidifies with pressure and one that activates dynamically (Fig. 1; Supplementary Table 1). Region I, which becomes more rigid with pressure, is defined by twenty-one methyl-bearing side chains that are largely localized to the N-terminal domain of the protein. Eight of these side chains are fully buried. Region II, which becomes more dynamic with applied pressure, is comprised of seventeen methyl-bearing side chains in the C-terminal domain and eleven of these are fully buried. Nine methyl-bearing side chains are outside of these regions. All methyl probes are 7 Å or more from the barnase-barstar interface, which is highly polar and extensively hydrated^25^.

A thermodynamic cycle from free barnase was created with barnase either bound to its inhibitor barstar, subjected to high hydrostatic pressure (3 kbar) or both (Fig. 1). At 3 kbar, binding of barnase to barstar results in an opposite response from the N-& C-terminal groups of side chains. Motion in Region I is activated by elevated pressure, which is opposite to the response of free barnase. Region II rigidifies upon barnase binding barstar, both at ambient and high pressure (Fig. 1; Supplementary Table 1). Application of high pressure to the barnase-barstar complex leads to a general increase in the internal motion of barnase, with the largest change centered in Region II. As might be expected, the pressure sensitivity of a side chain’s motion is reduced as the 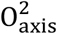 approaches the rigid limit of one at ambient pressure. Of the four states of barnase examined, the complexed state at ambient pressure is the most rigid 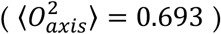 (Supplementary Table 1). The heterogeneous response of side chain dynamics to pressure is in stark contrast with other metrics examined. For example, fast backbone motions are generally suppressed in response to pressure and without apparent grouping to regions I & II (Table S1). The heterogeneity and complexity in the dynamical character of the protein is also not apparent from the more usual tactic of characterizing the pressure dependence of NMR chemical shifts and peak intensities (Supplementary Fig 2).

The localized response of motion to pressure is not easily explained by <O^2^_axis_> values at ambient pressure (Region I: <O^2^_axis_> (*var*O^2^_axis_) = 0.721 (0.031), n=16; Region II: <O^2^_axis_> (*var*O^2^_axis_) = 0.621 (0.030), n=12). Elevation to 3 kbar, results in <ΔO^2^_axis_> (*var*ΔO^2^_axis_) of +0.040 (0.002) and −0.053 (0.003) for Regions I and II, respectively. The effect of pressure is fundamentally related to changes in system volume. To examine potential contributions to the pressure sensitivity by the protein itself, we carried out a fine-grained volumetric analysis of the crystal structure (see Methods). Focusing on methyl-bearing side chains without solvent accessible surface area, we find Region I side chains in the ambient pressure structure, on average, have 35 Å^3^ more unoccupied volume surrounding the side chain than those of Region II (111 ± 23 Å^3^ and 78 ± 33 Å^3^, respectively; p < 0.022). Compression of voids explains the rigidification by pressure observed in Region I. In contrast, a more densely packed Region II may not be able to further compress and will respond to pressure through other mechanisms that decrease the system volume, such as local structural transitions or changes in hydration. These initial observations prompted us to compile a database of experimentally determined methyl symmetry axis order parameters and investigate the influence of surrounding void volume more generally.

The technical challenges of bridging crystallography and NMR relaxation are numerous. On the NMR side, relaxation datasets are sometimes compromised by contamination of relaxation from nearby nuclei, inadequate tumbling analysis, incomplete depositions, misassignments, and inclusion of data from ill-resolved resonances. Crystallography datasets often suffer from absent density maps, incorrect or incomplete modeling, low resolution, extreme experimental conditions (cryogenic temperatures), ambiguity in solvent and cosolvent identification, and poor representation of solvent and alternative conformations. There are approximately 2,568 experimentally determined O^2^_axis_ values of proteins with cryogenic crystal structures determined to high-resolution. We performed a fine-grained volumetric analysis of 42 crystal structures with corresponding 1027 measured 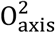 values derived from fully buried side chains that meet the quality criteria outlined in the methods section. All of these methyl-bearing side chains were observed to have surrounding void volumes less than 180 Å^3^. An intriguing trend reinforces the notion that there exists a lower limit on the 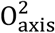 values imposed by the available surrounding unoccupied volume (Fig. 2). Side chains surrounded by less than 70 Å^3^ of void volume retain 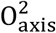 values above 0.5 with the large majority (75%) retaining high values above 0.75. Surrounding void volumes between 70 and 140 Å^3^ correlate with the entire range of possible 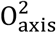 values. The range of void volumes over which low 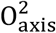 values become populated represents a window of about 50 Å^3^, which is a remarkably small fraction of the total unoccupied volume in proteins. Indeed, such a change in void volume is on the order of that made accessible by a single alternative rotamer or the removal of a single water molecule (Fig. 3; see text below).

**Figure 2.**
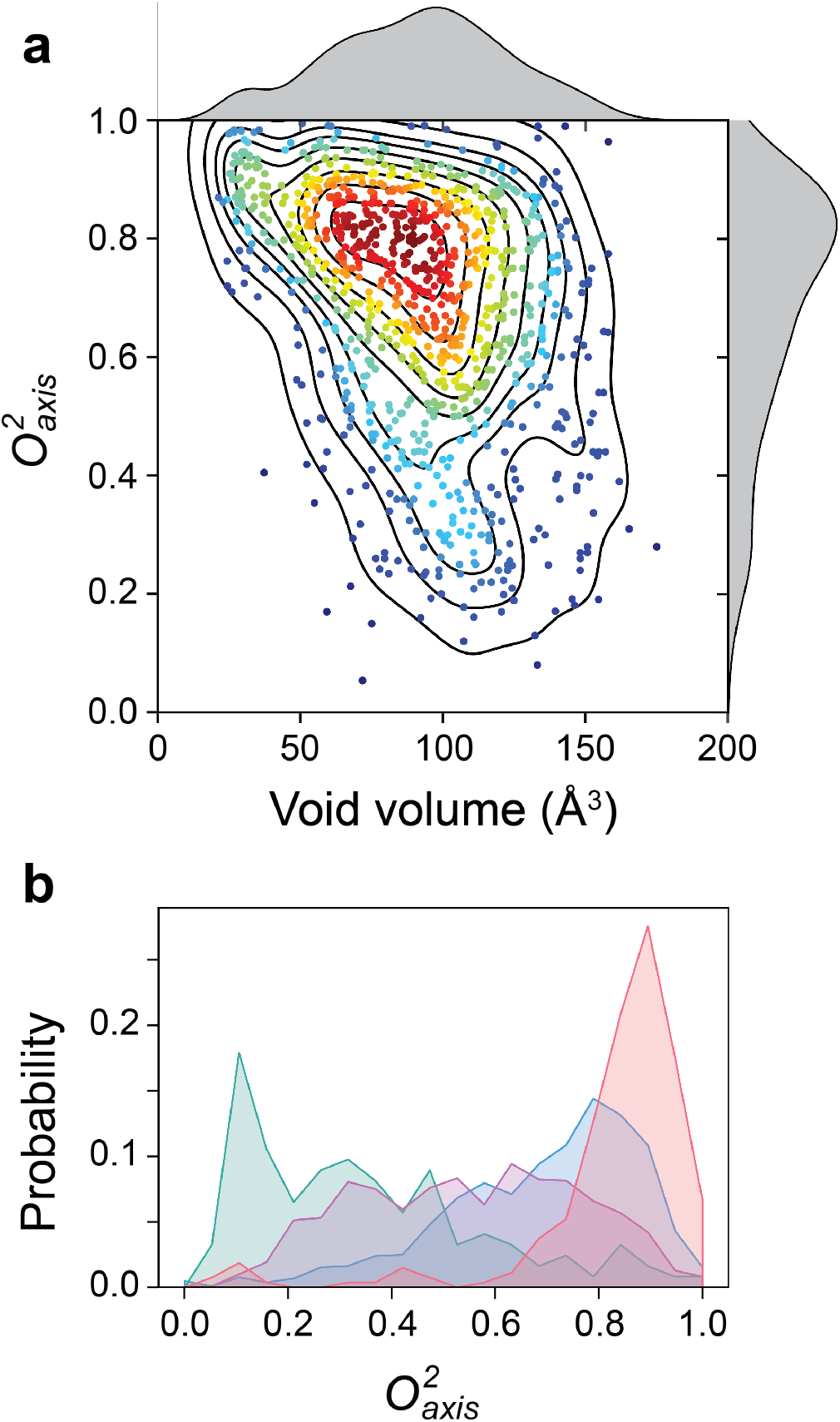
Dependence of fast side chain motion on void volume. **a**, Summary of the database of 50 cryogenic crystal structures and 2,568 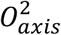 parameters obtained in solution near room temperature. Each filled circle represents one 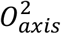 value plotted as a function of void volume calculated from the side chain’s corresponding crystal structure. Circles are colored from blue to red (sparse to dense) according to the contour lines. Sources are summarized in Supplementary Table 3. **b**, Distributions of experimentally determined O^2^_axis_ used in the analysis (see main text). The distributions for Nχ = 0 (Ala βCH_3_), Nχ = 1 (Thr, Val and Ile γCH_3_), Nχ = 2 (Leu and Ile δCH_3_), and Nχ = 3 (Met εCH_3_) were comprised of 273, 1059, 1111 and 125 values, respectively. Data sources are identified in Supplementary Table 1. [1 column]

**Fig. 3.**
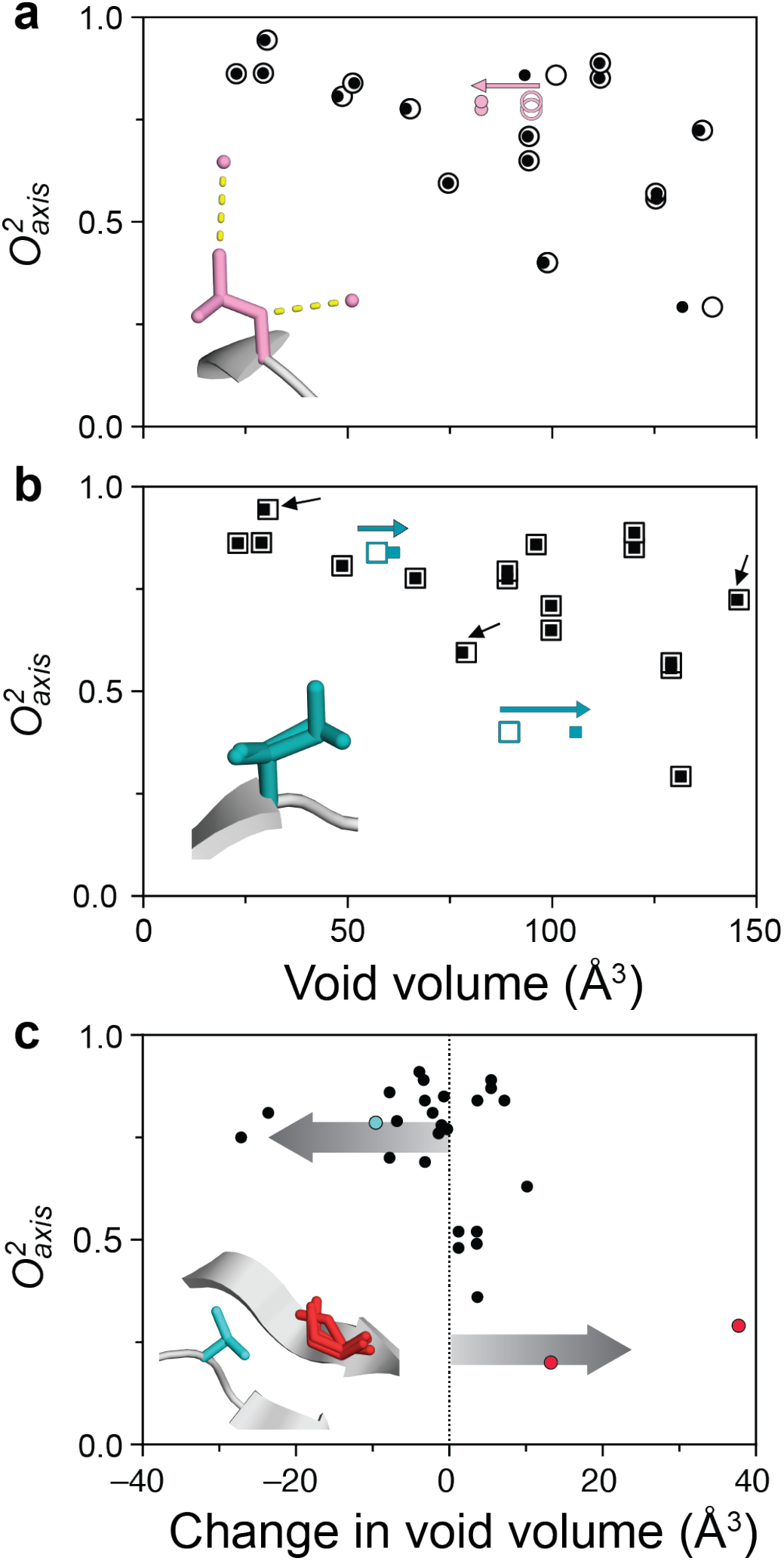
Relationship between surrounding unoccupied volume and fast methyl-bearing side chain motion. **a**, Methyl-bearing side chain 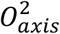 values correlated with the surrounding void volumes in the cryogenic structure of barnase (1B2X) calculated with (closed circles) and without (open circles) crystallographic water molecules. **b**, Void volumes of methyl groups in the room temperature structure of barnase (1A2P) with (closed squares) and without (open squares) the alternative conformation of Ile-96. **c**, The changes in void volume resulting from incorporation of a multi-conformational model of the room temperature structure of dihydrofolate reductase (4PST). See text. [1 column]

What defines 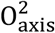 values above the identified lower limit of void volume clearly has a complicated origin. For example, many side chains with >100 Å^3^ of surrounding void volume can still display very high 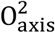 values. To identify the reasons for this behavior, we analyzed the role of three factors: the dependence on side chain topology (number of “soft” torsion angles); the presence of crystallographically defined solvent; and the presence of alternative side chain conformations. The distribution of 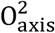 values (Fig 3a) reflects the long-appreciated feature that the *accessible* range depends on the number of torsion angles available in the context of a rigid backbone though the origin of the distribution within that range has remained a mystery^9,26^. It is also apparent that it is important to consider the presence of crystallographically defined solvent water molecules. For example, Leu-42 in barnase abuts two deeply buried water molecules, present in all 35 deposited crystal structures of barnase, which appreciably reduce the volume that would be otherwise available to the side chain (Fig 3b; 83 Å^3^ versus 95 Å^3^). Analysis of intermolecular dipolar relaxation from hydrogen-bonding partners (Trp-35 indole and Ile-51) to the two waters confirm their rigid presence in solution, characterized by long-residence times and restricted motion^4^. In general, underrepresentation of solvent molecules in crystal structures will result in overestimation of the void volume of surrounding side chains. Consideration of hydration water, which experience a broad range of residence times and motion while bound to the protein^27^, will likely be particularly important to understanding the influences of specific solvation.

Alternative conformations can also influence the effective void volume available. For example, including the alternative conformation observed for Ile-96 of barnase results in a large increase of its available volume, but also decreases the available volume of nearby side chains Ala-11, Leu-14, and Ile-88 (black arrows, Fig. 3c). The potentially large impact of alternative conformations is further illustrated by the multi-conformer model for the room temperature crystal structure of dihydrofolate reductase (DHFR)^28^. Including multiple side chain rotamers to represent the electron density results in an overall decrease of apparent void volume for the more rigid side chains and an increase for the more dynamic side chains (Fig. 3d). In other words, the underrepresentation of alternative conformations in cryogenic crystal structures leads to less accurate calculation of void volumes and deteriorates the correspondence with side chain dynamics. This implies that the broad distribution of low (<0.5) 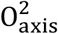 values with surrounding void volume (Fig. 2a) is likely due to inadequate representation of alternative conformations in single-conformer cryogenic crystal structures.

The remarkable sensitivity of protein conformational entropy to packing density has been predicted by simulation, albeit involving somewhat greater changes than identified here^29^. Indeed, analysis of hard-sphere liquids also indicates a dependence of dynamics on available free-space^30^ and reinforces the notion derived from the pressure sensitivity of fast aromatic ring motion that proteins interiors are quite liquid-like^31^. A heterogeneous response of protein domains to pressure was also reflected in the sole previous study of the pressure sensitivity of fast methyl-bearing side chain motion^32^. The influence of elevated hydrostatic pressure on the internal motion (conformational entropy) in the barnase-barstar complex, the first complex studied by these methods, is striking. High affinity binding selectively alters the response of barnase to pressure. Region I rigidifies as free barnase is compressed, indicating that there is conformational heterogeneity involving less efficiently packed alternative conformation(s). Interestingly, this region corresponds to a late-folding intermediate^33^. In contrast, pressure favors increased disorder on the sub-nanosecond timescale in Region II. Very localized spatial clustering of the response of fast motion to pressure was observed in ubiquitin^32^, but the degree of structural segregation in barnase is striking. The femtomolar affinity of the barnase-barstar complex exists despite a penalty by -TΔSconf of +11.7 kJ/mol. But at high pressure, the *overall* change in side chain dynamics is zero and binding occurs with no conformational entropy penalty. The response of side chains to pressure is consistent with an important role of conformational dynamics in the adaptation of protein function to extreme environments^23^. Furthermore, these results make clear that changes in both the magnitude and the sign of regional contributions of conformational entropy to the thermodynamics of protein function are possible. In a very real sense, the establishment of a connection between extremely subtle structural features (imperfect packing) and conformational entropy provides a mechanism for Nature to install the 35 year-old speculation of allostery without (detectable) structural change^34^.

## Methods

### Sample preparation

pET-DUET expression plasmids containing the genes for barnase and barstar under the control of their own T7 promoter were obtained from GenScript Biotech Corporation (Piscataway, NJ, USA). An N-terminal 6xHis-tag followed by a Factor Xa cleavage site (MGSSHHHHHHSQAPIEGR) was added to barnase while barstar remained untagged. Expression was carried out in BL21-(DE3) E. coli cells. Barstar expressed and purified with the N-terminal Met residue present. NMR relaxation samples were prepared largely as described elsewhere^4^. Deuterium and ^15^N relaxation experiments of the free proteins were performed on a 1:2 mixture of uniformly ^15^N-labeled protein and uniformly ^13^C-labeled protein expressed in 60% D_2_O media to generate the ^13^CH_2_D isotopomer. The complex was studied by combining ^15^N-labeled protein (barnase or barstar) with ^13^CH_2_D-labeled binding partner (barstar or barnase). Prochiral methyl assignments samples were expressed during growth on 10% ^13^C_6_-glucose and 90% unlabeled glucose and uniform ^15^N labeling^35^.

The barnase-barstar complex was isolated by Ni-NTA affinity chromatography and the complex dissociated with 6 M guanidine HCl, pH 7.9. Barstar was collected in the flow-through (20 mL) and refolded by dilution into 1 L of water. Refolded barstar was further purified by DEAE ion exchange column that included a wash with 25 mM imidazole, pH 7.9, 10 mM KCl and elution with 500 mM NaCl, spin-concentrated (3 kDa cutoff) and further purified by size exclusion chromatography on Superdex SEC-75 equilibrated with 25 mM imidazole, pH 7.9, 10 mM KCl. Barnase was eluted from the Ni-NTA column with 500 mM imidazole, spin-concentrated (3 kDa cutoff), and buffer exchanged into 25 mM imidazole, pH 6.2, 10 mM KCl and 5 mM CaCl_2_. The His-tag was cleaved with four µg of Factor Xa per mg of barnase added and mixed overnight at room temperature. The solution was passed through a 1 mL Ni-NTA column coupled to a SEC-75 in 50 mM imidazole, pH 7.9, 50 mM KCl, spin-concentrated and buffer exchanged to 25 mM imidazole, pH 6.2, 10 mM KCl. NMR experiments were performed with samples prepared in 25 mM imidazole, pH 6.2, 10 mM KCl, 5% D_2_O, and 0.02% NaN_3_ (w/v). Samples were stable about 1 month for the free proteins and several months for the complex at 35 °C.

### NMR assignment and relaxation of free and bound barnase

All experiments were carried out at 35 °C. Assignment experiments were done on a uniformly ^13^C, ^15^N-labeled sample, with only one protein in the complex labeled to reduce spectral crowding. Non-uniform sampling was used extensively for triple resonance assignment spectra^36^. Assignments were mapped to high pressure by collecting ^13^C and ^15^N HSQC spectra every 500 bar. These experiments were collected either at 500 or 600 MHz. Carbon and nitrogen HSQC spectra were collected at 1, 50, 500, 1000, 1500, 2000, 2500, and 3000 bar with a waiting period of 1h between each. Spectra were collected during ramp up and ramp down of pressure, with no detectable difference observed between them. Chemical shift analysis utilized the gyromagnetic ratio weighted change in chemical shift of ^1^H and ^15^N (or ^13^C) of bonded atoms resolved in two-dimensional correlation spectra.

Resonance peak heights and volumes were obtained using NMRFAM-SPARKY^37^. Only fully resolved cross peaks that fitted well to a Lorentzian lineshape to provide intensities were included. Peaks were integrated using the fitted Lorentzian function. The pressure dependence of gyromagnetic weighted chemical shifts were analyzed using a second-order Taylor expansion^38^. The correlation between second and first order coefficients was fitted with a linear regression with intercepts (6.5 ± 0.8) and (−16 ± 6) × 10^−10^ ppm bar^-2^, and slopes (−7.2 ± 0.2) and (−2.2 ± 0.2) × 10^−5^ bar^-1^ for amide proton and nitrogen, respectively, for free barnase, and intercepts (−2.2 ± 0.4) and (10 ± 2) × 10^−10^ ppm bar^-2^, and slopes (−6.7 ± 0.1) and (−3.2 ± 0.1) × 10^−5^ bar^-1^ for amide proton and nitrogen, respectively, for complexed barnase (Supplementary Fig. 1).

Longitudinal and transverse relaxation was measured using HSQC spectra with 9 interleaved delay points and 3 duplicates (delays 2, 5 and 8) for uncertainty estimation (each applied to itself and neighboring delay points 1-3, 4-6, and 7-9)^39^. Maximum peak intensities and uncertainties were used to fit single-exponential decay curves with 3 parameters. ^1^H-^15^N nuclear Overhauser enhancement (NOE) experiments were measured with a 5 s mixing time with and without irradiation of ^1^H. Relaxation was measured at 500 and 600 MHz (^1^H) for high-pressure experiments, and 500, 600, and 750 MHz (^1^H) for ambient pressure experiments. Deuterium relaxation employed Dz and Dy experiments with on-the-fly IzCz-compensation^40^. High pressure NMR relaxation experiments were carried out in a 3 kbar rated 5 mm o.d./3 mm i.d. ceramic NMR tube connected to a high-pressure Xtreme-60 pressure-stat syringe pump (Daedalus Innovations LLC, Aston, PA). The pressure medium was degassed water with a mineral oil interface with the sample. The effect of pressure on imidazole’s pK_a_ is small^41^. Relaxation measurements on the complex (24 kDa) were performed on uniformly ^15^N-labeled barstar and uniformly ^13^C-labeled barnase grown in 60% D_2_O.

The macromolecular rotational correlation model and Lipari-Szabo squared generalized order parameters of the amide N-H bond vectors 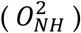 were determined from ^15^N relaxation experiments^42^. Tumbling models for the complex were obtained from ^15^N-labeled barstar backbone relaxation measurements. Tumbling models were chosen according to the Akaike and Bayesian information criteria and F-tests to obtain probability values for each model^43^. Simple model-free parameters were determined using a grid-search in a C++/AMP implementation of Relxn2A^4,44^. The analysis used an effective N-H bond length of 1.04 Å^26^, a general ^15^N tensor breadth of −170 p.p.m.^45^, a quadrupolar coupling constant of 167 kHz^46^, and a methyl rotation order parameter O^2^_rot_ of 0.1107 assuming perfect tetrahedral geometry of the methyl carbon used to extract the O^2^_axis_.

### Void volume calculations of protein structures

Voronoi volumes^47^ are ideally suited to investigate the volume of buried atoms as “the sum of polyhedral volumes is exactly equal to the total space occupied by the points”^48^. Voronoi volumes were determined with an in-house Cython program. Only structural models with a nominal resolution of < 2.5 Å based on data obtained at cryogenic temperature for the protein with the same biological context as that of the NMR experiment (e.g., free vs. complexed) were considered. Available room temperature crystal structures were analyzed separately. When multiple copies of the protein were present in the asymmetric unit, the copy with the strongest electron density and highest quality model was identified by visual inspection. The calculation finds the edges of the coordinates and defines a box with a 5 Å padding. A cubic grid is created with a step size of 0.01 Å. Each voxel is interrogated for the nearest heavy atom and assigned to it. Voxels within the van der Waals radius^49^ of any atom were excluded. The sum of all voxels assigned to an atom represents the atom’s Voronoi polyhedron and is used to calculate its volume. Side chain volumes were summed starting at the Cβ and ending with the atoms with the same number of dihedral angles as the methyl group of interest. For example, Ile Cγ2 methyls will include the volume of both Cγ carbons and one Cβ atom, while Ile Cδ methyls will include one Cδ, both Cγ, and one Cβ atom. The surface was defined using a large probe (2.4 Å radius) to avoid fitting inside any internal cavities. The surface algorithm ignored ligands and waters, so binding pockets were considered open surfaces as well. The probe was moved through the grid to find all voxels where the probe fit without steric overlap, and protein atoms that came within 1 voxel of the probe were flagged as belonging to the surface. If any atom of a side chain (starting at the Cβ and ignoring backbone) contained a surface atom, the side chain was not considered buried. Alternative rotamers were analyzed by including all rotamers in the calculation, summing the volumes of all rotamers of a given side chain, and subtracting the van der Waals volume only once. All unoccupied void volume is obtained by this calculation including that which remains from perfect packing of spheres.

## Data Availability

The barnase relaxation data reported here has been deposited to the BMRB under accession number @.

## Acknowledgements

We thank Kim Sharp and Ken Dill for helpful discussion. Funding from NIH (R01 GM102447), the University of Pennsylvania and Texas A&M University.

## Author contributions

A.J.W. and J.A.C conceived and designed the project. K.G.V. and J.A.C. carried out the relaxation experiments and data analysis. J.A.C. wrote the necessary software and carried out the volumetric analyses reported. J.A.C., K.G.V. and A.J.W. analyzed the results. A.J.W. and J.A.C. wrote the manuscript.

## Additional information

### Supplementary information

The online version contains supplementary material available at @ Correspondence and requests for materials should be addressed to A.J.W.

Reprints and permissions information is available at http://www.nature.com/reprints.

## Supplementary Information

**Supplementary Data Table 1:** Fast internal motion in barnase^a^.

| Barnase<sup>b</sup> | P (bar) | $\langle O_{axis}^2 \rangle$ | $var O_{axis}^2$<sup>d</sup> | n<sub>NH</sub><sup>e</sup> | $\langle O_{NH}^2 \rangle$ | $var O_{NH}^2$ | $\tau_m$ (ns) |
| --- | --- | --- | --- | --- | --- | --- | --- |
| Free (all)<sup>c</sup> | 1 | 0.653 | 0.046 | 96 | 0.851 | 0.002 | 4.78<sup>h</sup> |
| Free (Region I)<sup>f</sup> | 1 | 0.721 | 0.031 |  |  |  |  |
| Free (Region II)<sup>g</sup> | 1 | 0.621 | 0.030 |  |  |  |  |
| Free (all) | 3,000 | 0.647 | 0.054 | 92 | 0.866 | 0.001 | 5.94<sup>i</sup> |
| Free (Region I) | 3,000 | 0.762 | 0.029 |  |  |  |  |
| Free (Region II) | 3,000 | 0.568 | 0.039 |  |  |  |  |
| Complex (all) | 1 | 0.693 | 0.055 | 97 | 0.847 | 0.001 | 8.89<sup>h</sup> |
| Complex (Region I) | 1 | 0.750 | 0.030 |  |  |  |  |
| Complex (Region II) | 1 | 0.667 | 0.040 |  |  |  |  |
| Complex (all) | 3,000 | 0.649 | 0.051 | 90 | 0.899 | 0.001 | 11.48<sup>i</sup> |
| Complex (Region I) | 3,000 | 0.725 | 0.023 |  |  |  |  |
| Complex (Region II) | 3,000 | 0.589 | 0.048 |  |  |  |  |

<sup>a</sup> all data obtained at 308 K
<sup>b</sup> number of probes spectrally resolved in all four states are 33, 16, and 12 for all, Region I, and Region II, respectively.
<sup>c</sup> includes all spectrally resolved methyl probes in barnase
<sup>d</sup> variance is shown to indicate the variation of the dynamics within each region
<sup>e</sup> only amide <sup>15</sup>N-<sup>1</sup>H probes in barnase were used to characterize tumbling and backbone motion
<sup>f</sup> Region I includes the buried side chains of Val-10, Leu-14, Leu-42, and Ile-51 and the surface exposed side chains of Val-3, Ile-4, Thr-6, Leu-20, Thr-26, Ile-25, Leu-33, Val-45, and Thr-79
<sup>g</sup> Region II includes the buried side chains of Leu-63, Ile-76, Ile-88, Leu-89, Ile-96, and the surface exposed side chains of Leu-95, Thr-99, Thr-100, Thr-107, and Ile-109
<sup>h</sup> Fully anisotropic tumbling model, effective $\tau_m$ reported.
<sup>i</sup> Axially symmetric tumbling model, effective $\tau_m$ reported.

**Supplementary Table 2:**
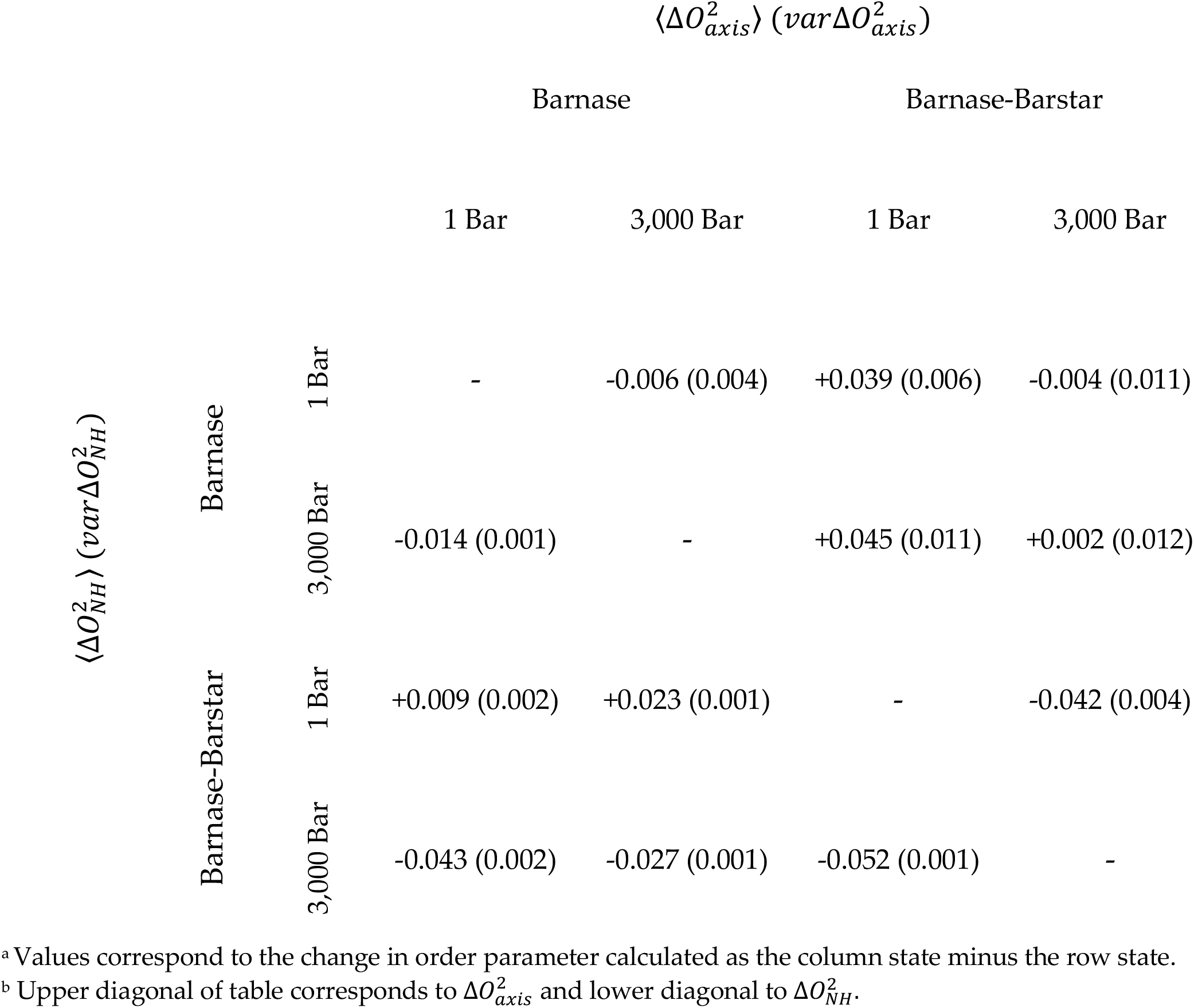
Changes in fast internal motion in barnase^a^.

**Supplementary Table 3:**
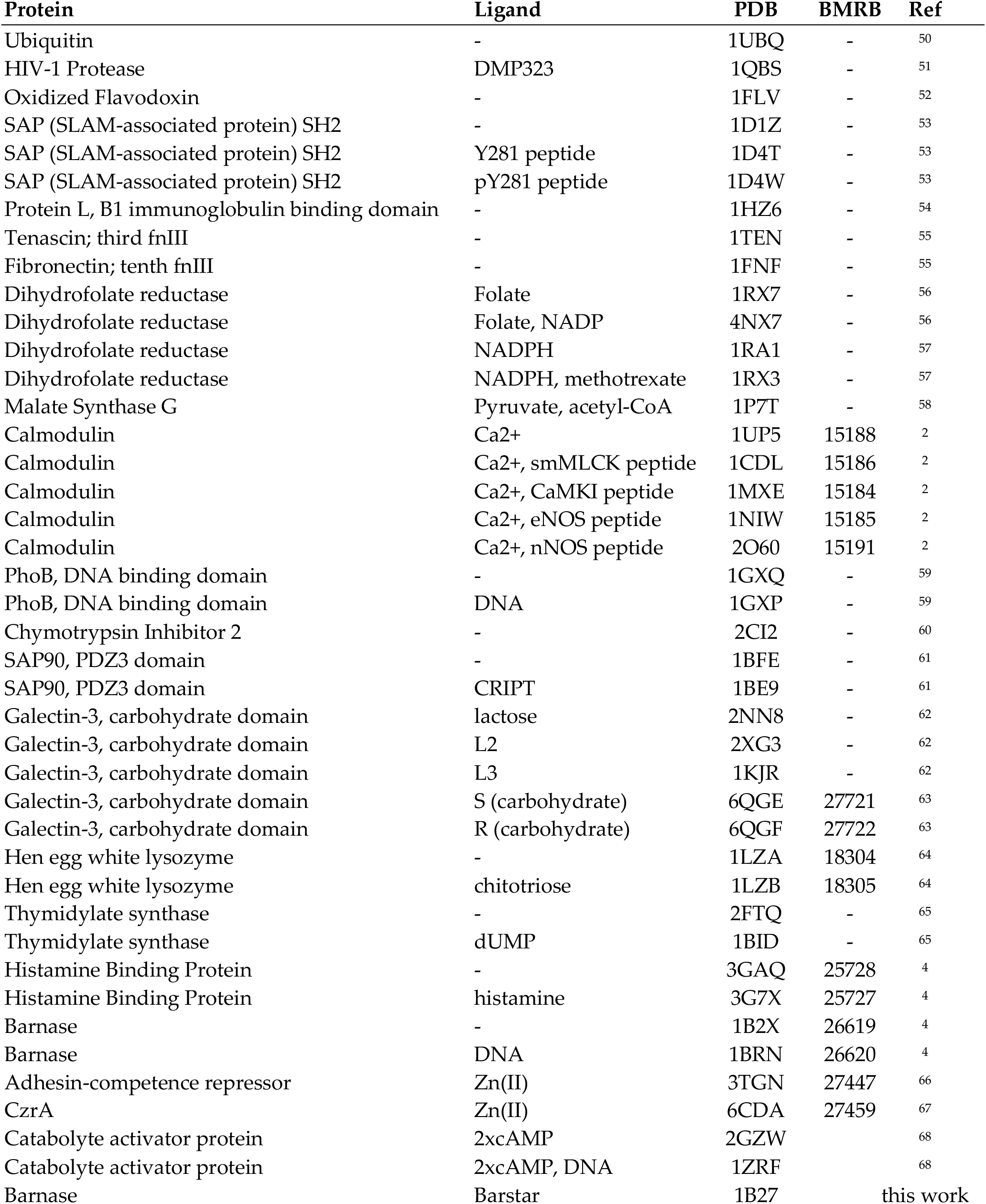
Summary of database used in analysis of surrounding unoccupied volume and local fast side chain motion.

**Supplementary Fig 1.**
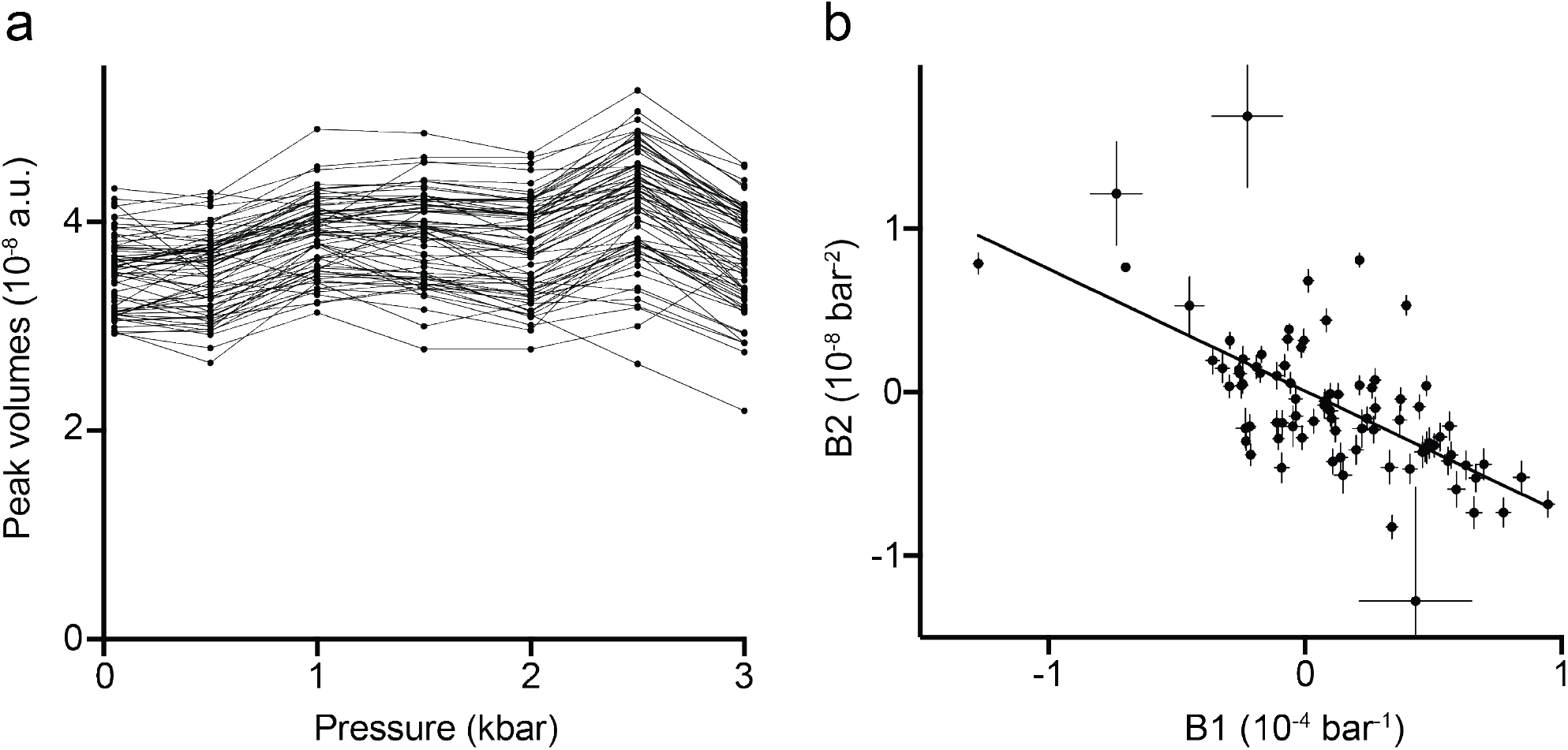
**a**, Dependence of free barnase ^1^H-^15^N HSQC resonance peak volumes on pressure. **b**, Correlation between first- and second- order terms of the polynomial used to fit ^1^H chemical shifts as a function of pressure. The line is a linear fit to the error-weighted data points and has an intercept of (6.5 ± 0.8) × 10^−10^ ppm bar^-2^ and a slope of (−7.2 ± 0.2) × 10^−5^ bar^-1^.

